# Interferon-stimulated genes in zebrafish and human define an ancient arsenal of antiviral immunity

**DOI:** 10.1101/693333

**Authors:** Jean-Pierre Levraud, Luc Jouneau, Valérie Briolat, Valerio Laghi, Pierre Boudinot

**Affiliations:** Institut Pasteur, Macrophages et Développement de l’Immunité, CNRS UMR3738, Paris, France; INRA, Virologie et Immunologie Moléculaires, Université Paris-Saclay, 78352 Jouy-en-Josas, France

## Abstract

The evolution of the interferon (IFN) system, the major innate antiviral mechanism of vertebrates, remains poorly understood. According to the detection of type I IFN genes in cartilaginous fish genomes, the system appeared 500My ago. However, the IFN system integrates many other components, most of which are encoded by IFN-stimulated genes (ISGs). To shed light on its evolution, we have used deep RNA sequencing to generate a comprehensive list of ISGs of zebrafish, taking advantage of the high quality genome annotation in this species. We analyzed larvae after inoculation of recombinant zebrafish type I IFN, or infection with chikungunya virus, a potent IFN inducer. We identified more than 400 zebrafish ISGs, defined as being either directly induced by IFN or induced by the virus in an IFN receptor-dependent manner. Their human orthologues were highly enriched in ISGs, particularly for highly-inducible genes. We identified 72 orthology groups containing ISGs in both zebrafish and human, revealing a core ancestral ISG repertoire, which includes most of the known signaling components of the IFN system. Many downstream effectors were also already present 450 My ago in the common ancestor of tetrapods and bony fish, and diversified as multi-gene families independently in the two lineages. A large proportion of the ISG repertoire is lineage-specific; around 40% of protein-coding zebrafish ISGs had no human orthologue. We identified 14 fish-specific gene families containing multiple ISGs, including finTRIMs. This work illuminates the evolution of the IFN system and provides a rich resource to explore new antiviral mechanisms.

**Key points:**

- We established an exhaustive list of larval zebrafish ISGs.
- Orthologous ISGs in fish and human identify a large ancestral ISG repertoire.

## Introduction

All living organisms are targeted by viruses, and evolution has given rise to various antiviral strategies. Vertebrates possess many unique immune features, including their principal innate antiviral system based on signaling by type I interferons (IFNs). Type I IFNs induce the expression of hundreds of proteins encoded by ISGs (IFN-stimulated genes), making cells refractory to viral infection (1). The origin of the IFN system, which seems to have replaced the RNA interference antiviral system still used by plants and most invertebrates (2, 3), is shrouded in mystery. Cartilaginous fish are the most basal clade with genomes containing recognizable type I IFN genes (4, 5), suggesting that this system appeared about 500My ago with jawed vertebrates. The IFN system integrates many components, most of which are encoded by ISGs, which can be traced back in genomes from distant clades. However, finding the orthologue(s) of a human ISG in another taxon does not imply that this gene is part of its IFN system. To understand the evolution of antiviral immunity, it is therefore desirable to establish how the repertoire of ISGs changed from early to modern vertebrates. This can be inferred by comparing the ISGs of current living representatives of distant vertebrate taxa.

Bony fishes (hereafter simply called “fish”) diverged from the tetrapod lineage about 450My ago, and, since viral infections are a major problem in aquaculture, their IFN system has been the subject of many studies, as reviewed in (6–9). Teleost fish possess several subgroups of type I IFNs (but no type III genes), with strong variation in gene numbers among fish taxa (8). The zebrafish possess four type I IFN genes, named *ifnphi1-4*; only *ifnphi1* and *ifnphi3* are active at the larval stage (10). Their receptors have been identified (10). Even before fish IFNs were known, the first fish ISGs were identified by homology cloning from cells stimulated by poly-I:C (11) or by differential transcript analysis of cells infected by viruses (12, 13). Because many virus-induced genes (vig) were homologous to well-known mammalian ISGs, they were hypothesized to be IFN-inducible, which was often confirmed by later studies, as in the case of *vig-1*, the *rsad2/viperin* orthologue (12, 14). Similarly, upon cloning of fish IFNs, induction of *Mx* (11) was used as a readout for their activity (15–17), confirming it was an ISG. The list of fish homologues of known ISGs rapidly grew with the release of new fish genomes and EST collections, allowing the development of micro-arrays to study fish response to virus or recombinant type I IFNs (18, 19). Candidate gene approaches were also developed, testing orthologues of known mammalian ISGs in qRT-PCR assays in multiple fish infection models (14, 20, 21). In parallel, approaches without a priori identified fish ISGs that had no orthologue in mammals, although they belonged to gene families involved in antiviral immunity. A large set of tripartite-motif protein-encoding genes, called *fintrims* (*ftr*), distantly related to *trim25* was identified in rainbow trout cells as a induced by virus infection (13) and later shown to form multigene families in teleosts, particularly extensive in zebrafish (22). Similarly, a family of IFN induced ADP-ribosyltransferases named *Gig2* was identified in crucian carp cells treated with UV-inactivated GCHV (grass carp hemorrhage virus) (23, 24). Some ISG were restricted to particular fish groups such as the non-coding *RNA vig2* that is found only in salmonids (25).

We previously established a list of zebrafish candidate ISGs using microarray analysis (26). For this, we compared the response to a poor IFN inducer, IHNV (infectious hematopoietic necrosis virus) (27) and a strong IFN inducer, CHIKV (chikungunya virus) (28). However, the array did not include the full complement of zebrafish genes, and the study identified virus-induced genes which were not necessarily ISGs. Here, to directly identify ISGs, we analyze the transcriptional response of zebrafish larvae injected with recombinant type I IFN. We rely on deep RNA sequencing, which is intrinsically quasi-exhaustive. Our approach is therefore limited mainly by the quality of genome assembly and annotation, which is excellent for the zebrafish (29). We complemented this analysis with a study of the response to CHIKV and its dependence to expression of the zebrafish IFN receptor chains *crfb1* and *crfb2* (10). We thus established a comprehensive list of ISGs of zebrafish larvae, and performed a detailed comparison with the human ISG repertoire. Our comparative analysis was facilitated by a compilation of human ISGs made to perform a systematic screen (30), and by the specialized database Interferome (31). We identify about 70 orthology groups that include ISG in both species and thus approximate the ISG repertoire of the common ancestor of all Osteichthyes. As ISGs typically evolve fast, with frequent duplications and gene loss, we also identify many families of fish-specific ISGs, which represent a rich resource for seeking new antiviral mechanisms. Our study provides a broad overview of the evolutionary patterns of genes belonging to the type I IFN pathway, and identifies gene modules induced by a viral infection independently of IFN.

## Materials and methods

### Zebrafish husbandry

Wild-type AB zebrafish, initially obtained from the Zebrafish International Resource Center (Eugene, OR, USA), were raised in the Institut Pasteur facility. Animal experiments were performed according to European Union guidelines for handling of laboratory animals (http://ec.europa.eu/environment/chemicals/lab_animals/home_en.htm) and were approved by the Institut Pasteur Animal Care and Use Committee. Eggs were obtained by marble-induced spawning, cleaned by treatment with 0.003% bleach for 5 minutes, and then kept in Petri dishes containing Volvic source water at 28°C. All timings in the text refer to the developmental stage at the reference temperature of 28.5°C. At 3 dpf (days post fertilization), shortly before injections, larvae that had not hatched spontaneously were manually dechorionated. Larvae were anesthetized with 200 μg/ml tricaine (A5040, Sigma-Aldrich) during the injection procedure.

### Interferon and virus inoculation

Recombinant zebrafish IFNφ1 (10), kindly provided by Rune Hartmann (University of Aarhus, Denmark), was inoculated by intravenous (IV) injection in the caudal cardinal vein. One nanoliter of 1mg/ml IFNφ1, or as a control, bovine serum albumin (BSA, New England Biolabs) in PBS1x/10% glycerol was injected. CHIKV infections were performed as described (26, 28). Briefly, ∼200PFU of CHIKV115 was injected IV in a volume of 1 nl at 3dpf.

### Interferon receptor knock-down

Morpholino antisense oligonucleotides (Gene Tools) were injected at the one to two cells stage as described (32). Two ng of *crfb1* splice morpholino (CGCCAAGATCATACCTGTAAAGTAA) was injected together with 2ng of *crfb2* splice morpholino (CTATGAATCCTCACCTAGGGTAAAC), knocking down all type I IFN receptors (10). Control morphants were injected with 4 ng of control morpholino (GAAAGCATGGCATCTGGATCATCGA) with no known target.

### RNA extraction

For RNAseq analysis, total RNA was extracted from replicate pools of 10 injected larvae (at 6 hours post injection for IFNφ1 treatment, or 24 hours post injection for CHIKV infections), using TRIzol (Invitrogen), following the manufacturer’s protocol. The integrity of the RNA was confirmed by laboratory-on-chip analysis using the 2100 Bioanalyzer (Agilent Technologies), using only samples with an RNA integrity number of at least 8.

### Illumina sequencing

Libraries were built using a Truseq mRNA-Seq Library Preparation Kit (Illumina, USA), according to the manufacturer’s recommendations. Quality control was performed on an Agilent Bioanalyzer. Sequencing was performed on a HiSeq 2500 system (Illumina, USA) and produced 65-base single-end reads.

### Mapping reads and gene expression counts

Sequences were trimmed using cutadapt (v1.8.3). The reads quality was checked with FastQC. Reads were then spliced-aligned to the zebrafish genome (GRCz10, Ensembl release 88) using TopHat2 (v2.0.14). The average number of mapped read per sample was 16.500.000. Only fragments mapping coherently and unambiguously to genes have been considered for gene counts. Gene counts have been assigned using featureCounts v1.5.2 (Liao, 2014).

### Identification of differentially expressed genes (DEG)

Differentially expressed nuclear genes between larvae treated with IFNφ1 and controls, between larvae infected by CHIKV and controls, or between *crfb1*+2 and control morphants all infected by CHIKV, were identified. DEG were identified using DESeq 1.18.0 (BioConductor) (Love et al., 2014) and R: 3–1-2 (R core team, 2017). Briefly, raw counts of genes were subjected to a minimal pre-filtering step: genes for which the count sum, per group of samples, was equal or higher than 10, in at least one group, were kept. Raw counts were normalized for library size and normalized data were fitted using a negative binomial general linear model. Data were adjusted for multiple testing using the Benjamini-Hochberg procedure (adjusted p value). Genes with an adjusted p value less than 0.01 and an absolute Fold Change (FC) > 2 or FC < 0.5 were considered as DEGs.

Sequence data were registered in the BioProject ncbi database (https://www.ncbi.nlm.nih.gov/bioproject) with the SRA accession number: PRJNA531581.

### Identification of human orthologues and ISGs

Orthology analysis was primarily based on data from the Ensembl database (www.ensembl.org), the zfin database (zfin.org), and the literature, notably our previous analysis of zebrafish orthologues of human ISGs (26). Data were systematically curated manually, and conflicts resolved using a combination of literature search, synteny analysis, and sequence homology analysis (two-way protein BLAST). When human genes were present on the list compiled by Schoggins et al. (30), they were labelled as ISGs. If absent from the list, gene names were further queried on the Interferome website (http://www.interferome.org/interferome/search/showSearch.jspx) which compiles the results of many transcriptomic studies on human and mouse samples after IFN stimulation (31). We postulated that human genes present in Interferome could be considered as ISG when being significantly induced at least 2-fold with a stimulation for no more 12 hours by type I IFN in at least 4 datasets.

### qRTPCR

RNA was extracted from individual larvae using RNeasy® Mini Kit (Qiagen). cDNA was obtained using M-MLV H- reverse-transcriptase (Promega) with a dT17 primer. Quantitative PCR was then performed on an ABI7300 thermocycler (Applied Biosystems) using Takyon™ ROX SYBR© 2x MasterMix (Eurogentec) in a final volume of 25 µl. The following pairs of primers were used: *ef1a* (housekeeping gene used for normalization): 5’-GCTGATCGTTGGAGTCAACA-3’ and 5’-ACAGACTTGACCTCAGTGGT-3’; mxa: 5’-GACCGTCTCTGATGTGGTTA-3’ and 5’-GCATGCTTTAGACTCTGGCT-3’; *ddx58*: 5’-ACGCCGGAGAAAGAATTTTTC-3’ and 5’-TCGACAGACTCTCGATGTTG-3’; *aqp9a*: 5’-CTGTACTACGACGCCTTCAT-3’ and 5’-GAGAATACAGAGCACCAGCA-3’.

## Results

### RNAseq analysis of IFN*φ*1-regulated genes

To make an inventory of zebrafish ISGs, we first injected 3dpf larvae with recombinant zebrafish IFNφ1 the first type I IFN to be identified in zebrafish, or BSA as a negative control. Based on preliminary kinetic experiments, we chose 6 hours post injection as the early plateau phase of ISG expression (Figure S1A). RNA was extracted from multiple pools of ten larvae and subjected to deep sequencing using an Illumina-based platform sequencing. Reads were mapped to zebrafish genome (zv10), and the differential analysis performed using the DESeq package.

Choosing as cutoff values adjusted *p* values <5% and fold-change (FC) >2, we identified 360 IFNφ1 up-regulated genes (which are ISGs, by definition) and 75 down-regulated genes (Table S1). As expected, genes with high basal expression levels tended to display lower fold-change (Figure 1A). The top IFNφ1-upregulated genes (with FC >100) comprised many well-known ISGs, many previously used as IFN signature genes in zebrafish, including several *mx* genes, *rsad2, cmpk2*, several *ifit* genes, the ubiquitin-like *isg15*, the helicase *dhx58* (aka *lgp2*), the kinase *pkz*, the transcription factor stat1b, and the chemokine ccl19a.2 (Figure 1B). To our surprise, *ddx58* (encoding RIG-I), a well-known and conserved ISG (33), was not found in that list. In fact, the gene model is missing on the zebrafish reference genome, with only fragments of the sequence present in the assembly. Therefore, we performed qRT-PCR for ddx58 and confirmed that it is induced by IFNϕ (Figure S1B). We searched the list of zebrafish orthologues of human ISGs (26) for more genes not annotated on the reference genome and found only one besides ddx58: aqp9a. By qRT-PCR, this gene appeared to be moderately (0.75 fold) downregulated by IFNϕ (Figure S1B), and thus was not an ISG.

**Figure 1.**
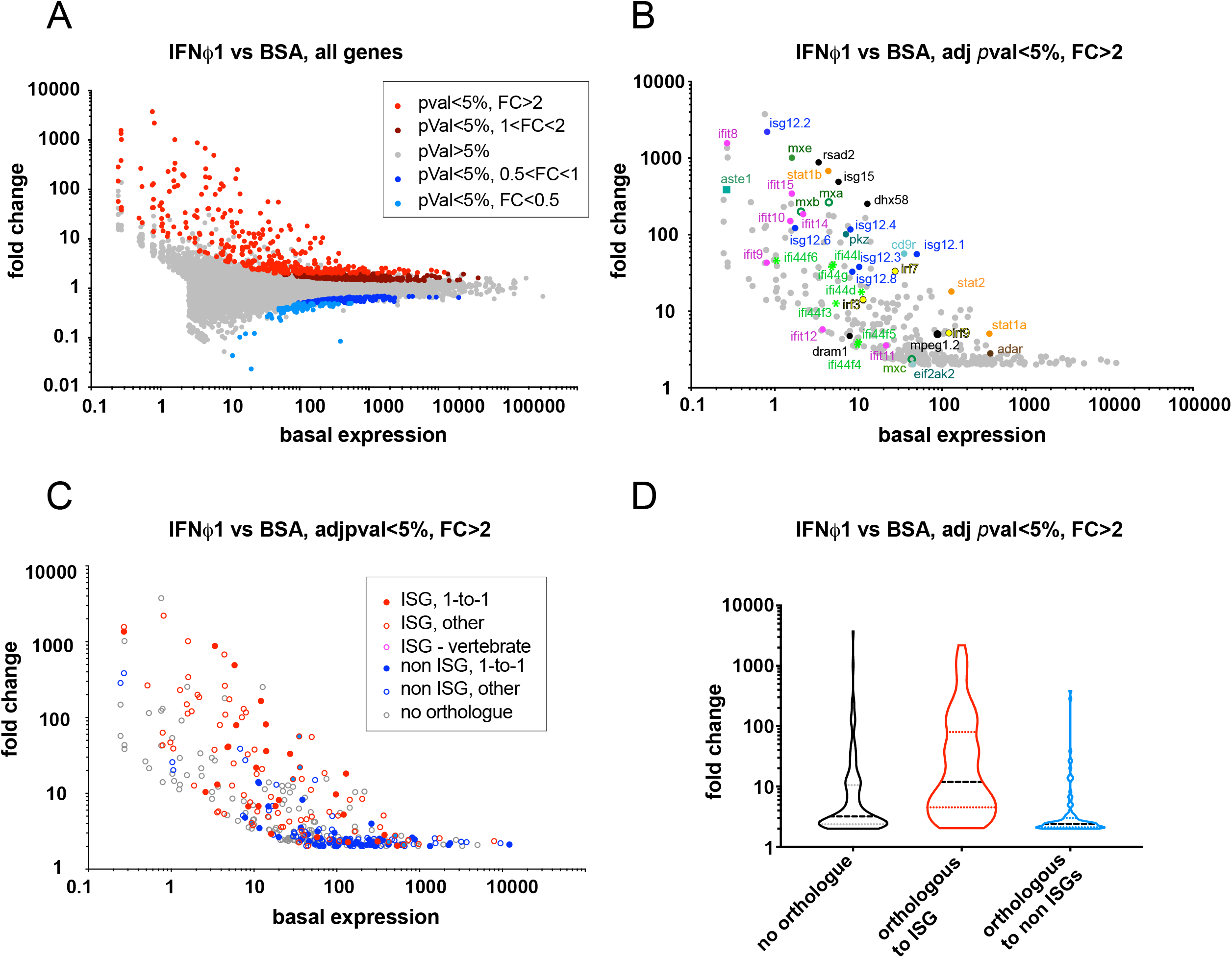
Zebrafish larva transcriptional response to type I IFN. (A) Fold change (FC)/basal expression representation with all genes detected in the analysis. Basal expression was the average of read numbers mapped onto a given gene in control (BSA-injected) larvae. FC is φ1-injected larvae divided by basal expression. pVal corresponds to adjusted p-Value. (B) FC/basal expression representation, limited to ISGs identified in IFNφ1 larvae (e.g. red dots in panel A); key genes commented in the text are indicated with gene families identified by different colors/symbols. (C) FC/basal expression representation for zebrafish ISGs represented according to the type of orthology with human genes, and if human orthologues include at least one human ISG. 1-to-1: single zebrafish gene orthologous to single human gene. Other: other orthology relationships between zebrafish and human genes (many-to-many, 1-to-many, many-to-1). Vertebrate: zebrafish gene sharing a common ancestor with a human ISG at the basal or jawed vertebrate level, but not orthologous. No orthologue: no orthology relationship with a human gene, and no known common ancestor with human ISG at vertebrate level. Non-ISG: orthologous human gene(s) do not include any ISG. (D) Fold change distribution for zebrafish ISG with different types of orthology relationship to human genes, using the same color code as in C but pooling genes with single and multiple human orthologues. Zebrafish ISGs with a human ISG within a vertebrate-level orthology groupes were not included in this analysis.

**Table 1.**
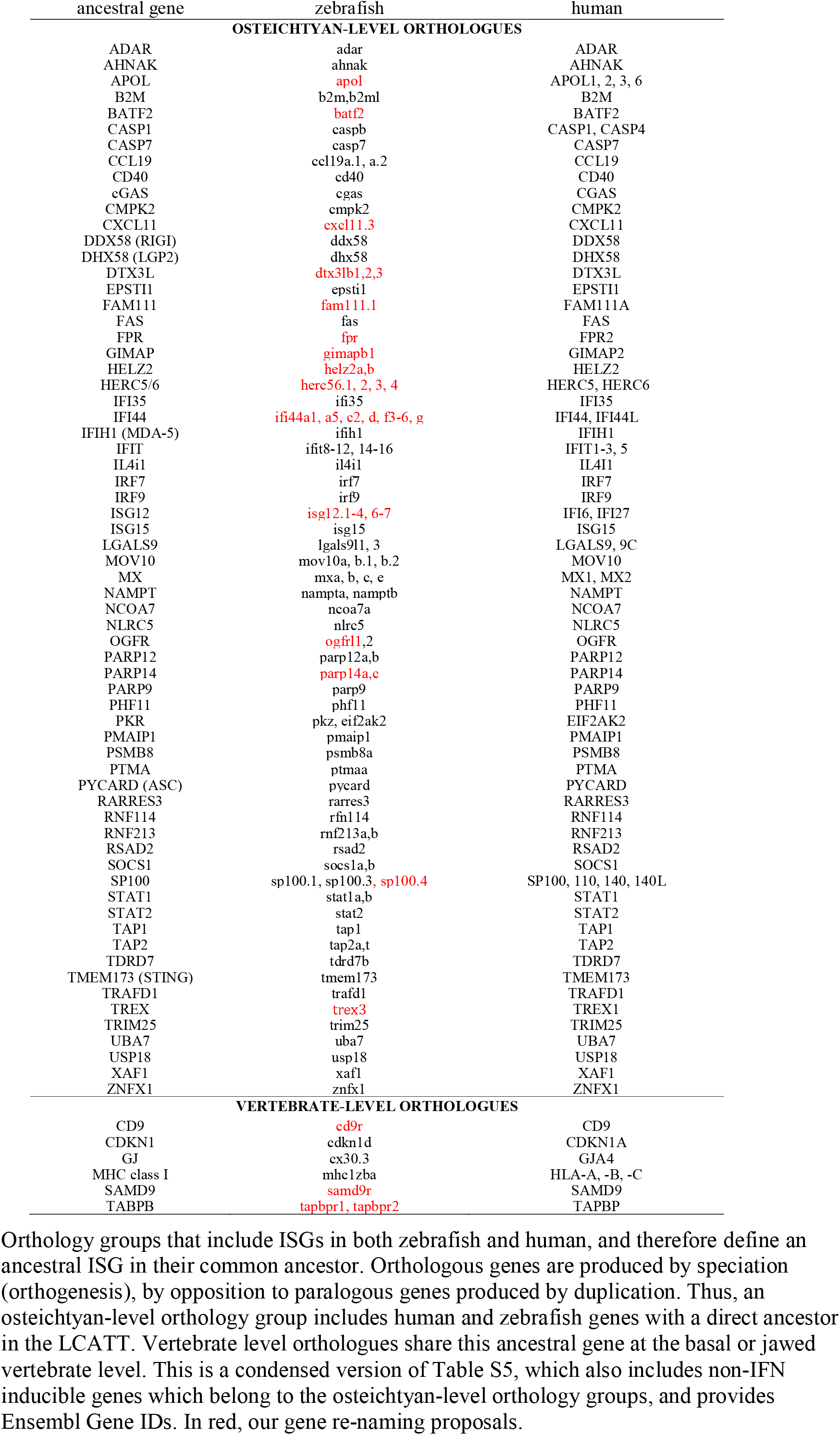
Orthology groups defining ancestral ISGs.

Among the 360 zebrafish ISGs identified by RNAseq, 23 corresponded to non-coding ISGs or transposons with no clear homologues in mammals, and were excluded from further phylogenetic analyses.

Gene ontology (GO) analysis was performed using David and Gorilla, and showed, as expected, that IFNφ1 upregulated genes were strongly enriched in genes linked to “antiviral response”, “type I interferon pathway” and “antigen processing and presentation” (not shown). Similarly, enrichment analyses identified KEGG pathways for Influenza, Measles and Herpes simplex infection as well as RIG-I signaling and cytosolic DNA sensing pathways. The list of downregulated genes was not found associated with any particularly notable function in these analyses.

### Human orthologues of zebrafish ISGs are enriched in ISGs

We then searched for the human orthologues of the 337 identified protein-coding zebrafish ISGs (Table S2). All types of orthology relationships between zebrafish and human were observed, from none to “many-to-many”. One-to-one orthology was found for 77 genes (Figure 1C). We identified one or several human orthologues for 200 zebrafish ISGs. This proportion (200/337, 59%) is significantly lower than the 71% reported for the whole genome (29) (Fisher’s exact test, p<0.0001).

We then searched which of these human genes were themselves ISGs. We found 61 ISGs present in the list of 446 human ISGs compiled by Schoggins et al. (30); by querying the Interferome database (31), we identified 11 additional human ISGs (Table S2). In total, 97 zebrafish IFNϕ genes were orthologous to at least one human ISGs (Figure 1C). In addition, we identified a handful of genes that were not true orthologues, but shared ancestry with a human ISG at the vertebrate level, such as MHC class I genes (see comments on Table S2).

As expected, human orthologues of zebrafish ISGs were strongly enriched in ISGs: while there are 446 human ISGs out of 20454 genes in the genome (i.e., 2%), we found 72 ISGs among the 196 human orthologues to zebrafish ISGs (i.e., 37%) (Fisher’s exact test, p< 0.0001). Interestingly, FC values of zebrafish ISGs were higher when they were orthologous to a human ISG than when their orthologues were not ISGs (red vs blue on Figure 1C-D), while FCs of zebrafish ISGs without human orthologues (grey on Figure 1C-D) were intermediate (mean values, 127.7, 10.6 and 46.9, respectively; all groups significantly different from each other, p<0.001, Kruskal-Wallis test). Thus, inducibility by type I IFNs is often evolutionary conserved, making it possible to infer an ancestral set of ISGs.

### Fish-specific ISG genes and families

Consistent with the expected high rate of duplication and divergence of ISGs, a significant proportion of zebrafish IFNφ1-up-regulated protein-coding genes had no identifiable orthologue in the human genome (137/337; 41%). Interestingly, but not unlike genes with human orthologues, many of these genes belonged to multigenic families. We show on Figure 2 the fish-specific gene families that contain several zebrafish ISGs, with domains identified by the SMART tool (34).

**Figure 2.**
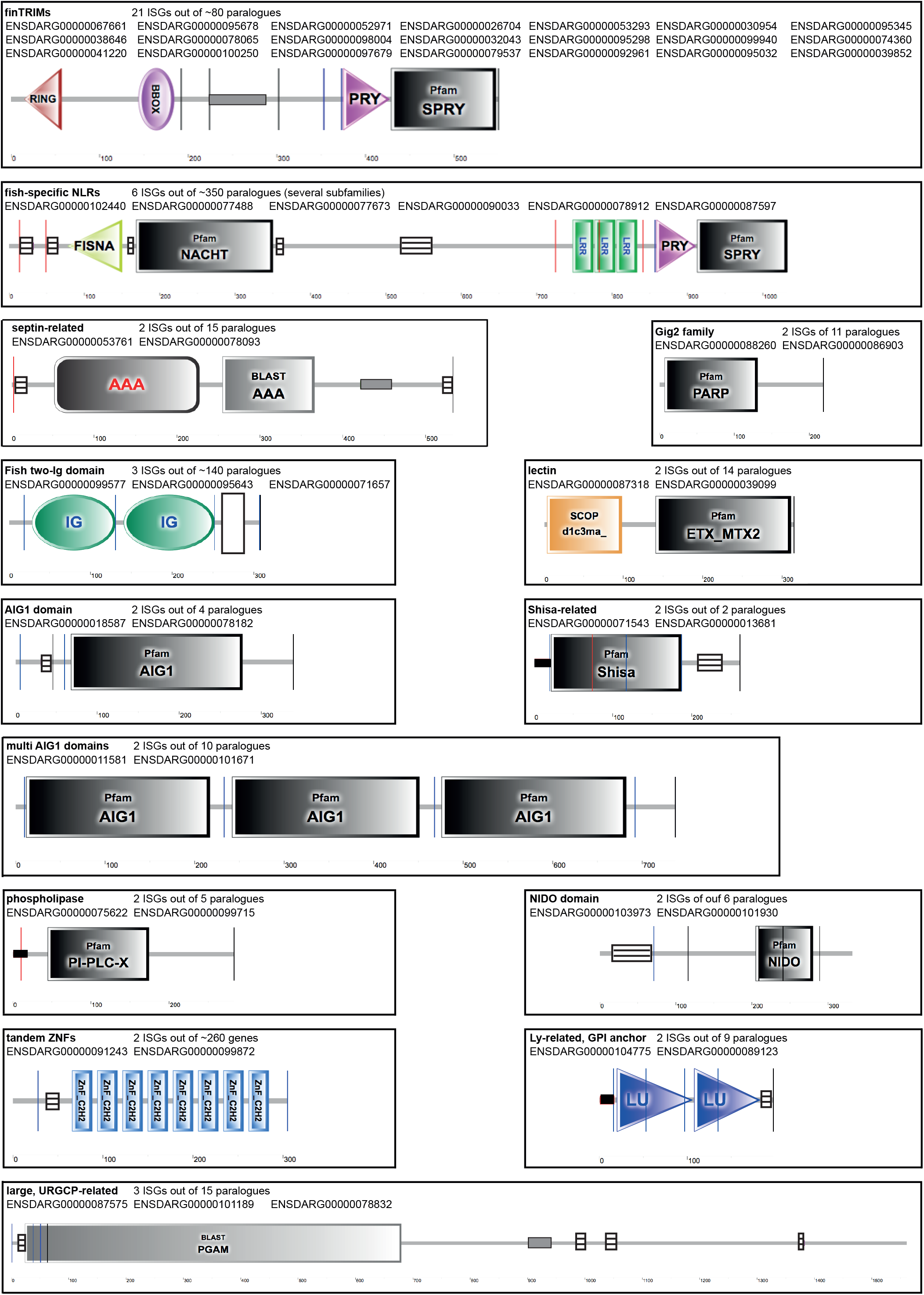
Main families of fish-specific ISG and their domain organisation. Each panel show the typical domain organization of a family of fish-specific ISGs (e.g., with no human orthologue), as determined by SMART analysis (http://smart.embl-heidelberg.de/). The accession numbers of IFNφ1 genes (FC>2, pVal<5%) and the fraction they represent within the family are indicated. Vertical lines represent exon boudaries. Besides named domains, boxes represent coiled-coil regions (grey), low complexity regsions (striped), transmembrane domains (black), and leader peptides (white).

The genes listed on Figure 2 included more than 20 *fintrim* (ftr) genes, a family identified first as virus-inducible genes in rainbow trout (13), highly diversified in zebrafish, and hypothesized to antagonize retroviruses (22). Of note, besides finTRIM, there are two other large TRIM gene expansions in zebrafish, each with a single human orthologue of unknown function (35). The btr (bloodthirsty-like TRIMs), related to human TRIM39, include several ISGs. By contrast, no member of the TRIM35-like family was upregulated by IFNφ1.

Another family had been described as virus-inducible in fish: *gig2* (GCRV-induced gene 2) genes (23). gig2 genes are distantly related to the PARP family (24), which include several ISGs in humans and zebrafish. The zebrafish *gig2p* and *gig2o* genes are induced by IFNφ1.

To our knowledge, the genes in the remaining fish-specific families had not previously been described to be interferon- or virus-inducible. These families are diverse, encoding proteins expected to be membrane receptors, and presumably secreted, nuclear, or cytosolic proteins (Figure 2).

Eight members of the very large NLR family were ISGs. These genes belong to groups 1, 2, 3 and 4 as defined in (36), and two of them belong to a fish specific subset defined by the presence of a C-terminal B30.2 (or PRY-SPRY) domain (37, 38), which is most similar to the corresponding domain of finTRIM genes (22). The specific function(s) of zebrafish NLR genes remain poorly understood, but this highly expanded family may be central for inflammatory mechanisms.

Additionally, 3 ISGs corresponded to membrane proteins with two immunoglobulin (Ig) domains and a transmembrane region but not ITAM or ITIM (immunoreceptor tyrosine-based activating/inhibiting motif). These genes belong to a very large family with 140 members, which we propose to name f2Ig (fish genes with two Ig domains).

### RNAseq analysis of CHIKV-induced genes

Experimental infection of zebrafish larvae with chikungunya virus (CHIKV) induces a strong type I IFN response (28). Our previous microarray-based analysis indicated that the response to CHIKV was dominated by ISGs (26). However, to allow comparison with another virus with slower IFN induction kinetics, this analysis had been performed at 48 hours post infection (hpi), while the peak of the IFN and ISG response, as determined by qRT-PCR, is at 24 hpi (26). Therefore, we re-analyzed here the transcriptome of CHIKV-infected larvae at 24 hpi using deep RNA sequencing. Choosing the same cutoff values as for the IFNφ1 analysis, we identified 466 CHIKV up-regulated genes and 26 down-regulated genes (Table S3). Hundreds of new CHIKV-inducible genes were identified, either because they were absent from the microarray, or because their induction were below the cutoff of the first analysis. Among the genes significantly upregulated in the microarray study, all those with a human ISG orthologue were also upregulated in this new dataset, and, as expected, typically much more (Figure S2).

About half of genes induced by IFNφ1 were also induced by the viral infection (181/360 Figure 3A; Table S3, yellow), including almost all (84 out of 97) genes orthologous to a human ISG, such as *mxa, b* and *e, stat1a and b; stat2; rasd2, isg15*, etc. There was a clear correlation of the FC values for genes induced by both IFNϕ and CHIKV (Figure 3B). However, almost two thirds of the genes induced by CHIKV infection were not significantly modulated by IFNφ1 (285/466; Figure 3A). We then asked whether this CHIKV-specific response could correspond essentially to genes for which there was weak induction by recombinant IFNφ, below our arbitrary cutoff. We therefore extracted from this list genes induced by IFNφ1 with FC>1.5 and with an adjusted *p* value<20%, and we found 66 genes matching these conditions (Table S3, green): 21 genes without annotation and 45 annotated genes, many of which were notoriously linked to the type I IFN system. These genes notably comprised *crfb1*, encoding a type I IFN receptor subunit, and two other cytokine receptors *il10Ra* and *il13Ra*; four chemokines (*ccl34, cxcl11.6, cxc18b and cxc20*); ten additional *fintrim* and 3 other members of the gig2 family; and two irf transcription factors (2 and 10). It also includes the metalloreductase steap3, whose mammalian orthologue is not an ISG, but regulates type I IFN response, CXCL10 induction and iron homeostasis in mouse macrophages (39).

**Figure 3.**
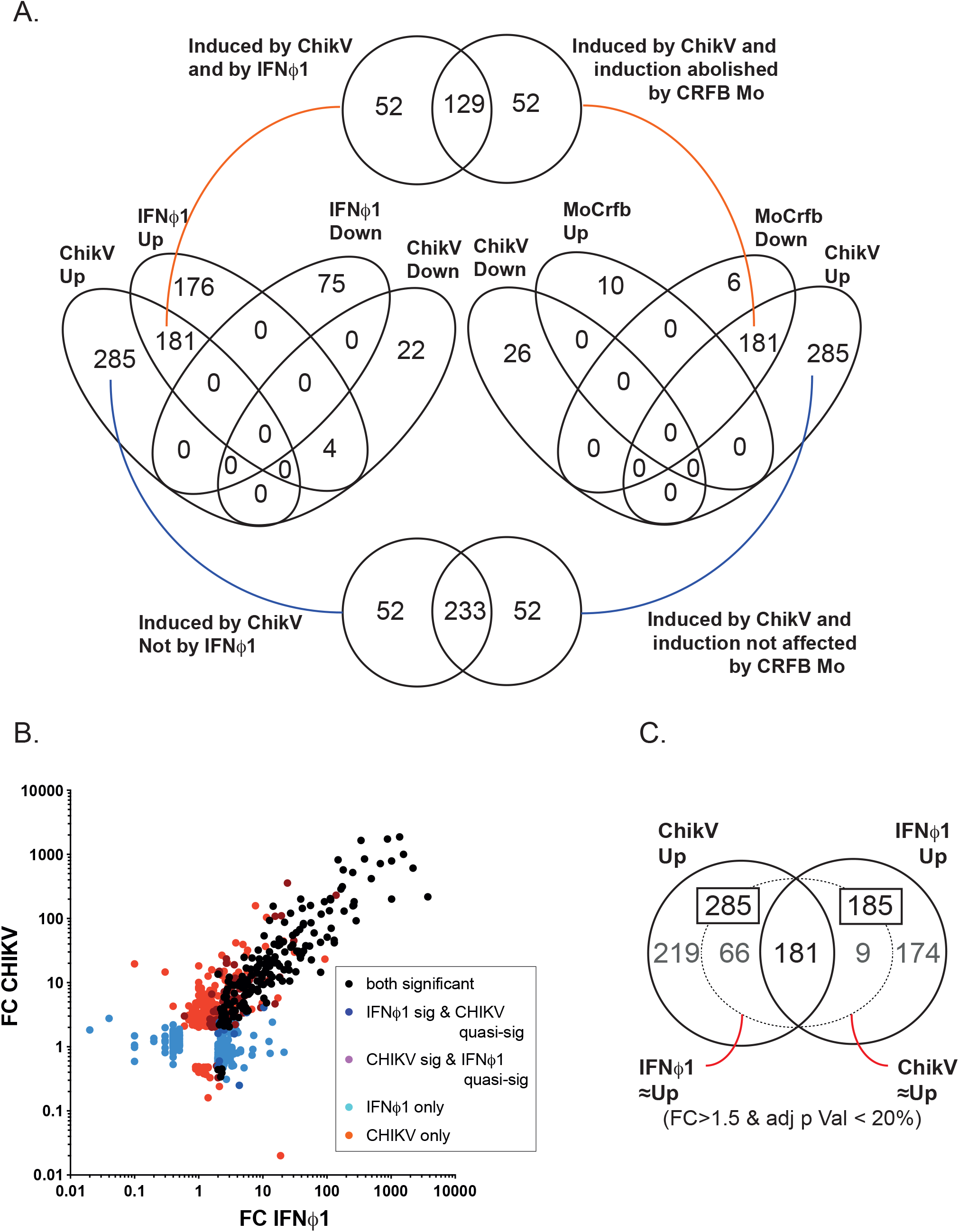
Comparison of ISG repertoires identified by IFNφ1 induction and by CHIKV infection. (A) Venn analysis of genes up-or down-regulated in zebrafish larva by IFNφ1 treatment (IFNφ1), CHIKV infection (ChikV), and CHIKV infection in the context of type I IFN receptor knock-down (“MoCrfb”). ISG are identified either by their responsiveness to IFNφ1 or by a CHIKV induction abolished in *crfb1*+2 morphants (venn diagram at the top). Genes induced by CHIKV infection in an IFNφ1 independent way are analysed in the Venn diagram at the bottom of the panel. (B) FC/FC representation of transcriptome response to IFNφ1 and CHIKV. Color code identifies genes significantly induced by IFNφ1 and/or CHIKV, and genes which are significantly induced in one condition (FC>2, adj p val< 0.05) while almost significantly induced in the other (“quasi Sig”; thresholds FC>1.5 and/or adj p value < 0.2). (C) Venn analysis of genes significantly induced by IFNφ1 and/or CHIKV. Gene subsets corresponding to genes induced quasi-significantly is represented within dotted lines.

Besides this intermediate gene set, a conservative list of 219 genes seems to be up-regulated only by the virus (FC>2 and adjusted p value <5%), independently of IFNφ1 (FC<1.5 or adjusted p value >20%) (Table S3, blue; Figure 3C). This list contains 105 genes without annotation, but also several functional modules providing interesting insights on the virus-host interactions. Functional analysis using DAVID identified 6 enriched KEGG pathways, namely Cytokine-cytokine receptor interaction, Cytosolic DNA-sensing, Toll-like receptor signaling, RIG-I-like receptor signaling, Proteasome, Herpes simplex infection.

Importantly, type I IFNs were induced by the infection. Consistent with our previous report with another virus (10), *ifnphi1*and *ifnphi3*were clearly dominant at this larval stage, with 58±3 and 47±11 reads respectively, compared with 9±4 reads for ifnphi2, and none detected for ifnphi4. Two pro-inflammatory cytokines *il1b* and *tnfb* were also upregulated. Among typical sensors, tlr3, mb21d1 (encoding cGAS) and its downstream adaptor tmem173 (encoding STING), and several kinases of the IFN signaling pathways (ripk1, tbk1) were present. Seven proteasome subunits are induced by the virus, suggesting activation of protein degradation and Ag presentation pathways. The complement pathway also stands out as an important module upregulated by CHIKV infection: twelve complement component genes (*c1, c2*, several *c3, c7; c9; cfB; cfhl-1,-3 and −5*) were induced by CHIKV, suggesting that it is an important defense triggered in a Type I IFN independent manner. Additionally, this response comprises 3 metalloaminopeptidases (anpepb, erap1b and 2); the myeloid markers *ncf1*, *mpx* and marco; 2 guanylate binding proteins (*gbp1* and *2*) that have well known orthologues in human; the transcription factors *atf3* and *irf1b*, and with a high level of expression, the enzyme *rnasel3* (an orthologue of human RNASE4, not of RNASEL, an ISG with no fish counterpart). Nine *fintrim* and 3 btr can also be noted, underscoring the importance of these TRIM with PRY/SPRY domains in virus host interactions altogether. Thus, CHIKV induces a typical IFN-stimulated response of high magnitude, but also a broader and less overt inflammatory response.

### IFN receptor dependence of the response to CHIKV

To test the IFN-dependence of the response to CHIKV, we used morpholinos to knock down in zebrafish larvae *crfb1* and *crfb2* which encode specific chains of the two types of type I IFN receptors of zebrafish (10). We previously showed that such IFNR morphant larvae are hypersusceptible to CHIKV infection, dying 2 to 3 days after virus injection (28). We analyzed by deep RNAseq their transcriptional response to CHIKV at 24 hours post-inoculation, and compared it to that of control morphant larvae. Choosing as cutoff values adjusted p values <5% and a ratio between IFN-R morphants and controls >2, we identified 187 genes for which induction was dampened by IFNR knockdown, and 10 genes that were upregulated in morphants (Table S4). Among CHIKV-induced genes (Table S3), 181 were IFNR-dependent, representing a significant fraction (181/466; 39%) (Figure 3A). Predictably, the list of genes upregulated by CHIKV in a IFNR-dependent manner largely - but not fully - overlapped with the gene set induced by recombinant IFNφ1 (129/181; 71%, see Figure 3A). This approach led us to classify 52 new zebrafish genes as ISGs, being induced by CHIKV in an IFNR-dependent manner, even if they were not significantly induced by recombinant IFNφ1. As previously, we searched the human orthologues of these additional ISGs (Table S2, bottom), identifying a few more human ISGs in this list, such as cGAS, NLRC5 or IFI35.

Together, our results provide a near-exhaustive list of zebrafish ISGs at the larval stage, identified by two independent approaches, and a useful reference for future studies.

### Ancestral ISGs

Assuming that the common ancestor of genes that are IFN-inducible in both human and zebrafish was itself an ISG in their last common ancestor ∼450 My ago, we can define a list of ancestral ISGs. We identified 66 orthology groups that included an ISG on both the human and the zebrafish sides (Table I, Table S5). A few more ancestral ISGs were also defined by pairs of ISGs with orthology relationships at the early vertebrate or gnathostome level – meaning that the zebrafish gene is not directly orthologous to a human ISG, but is paralogous (with an ancestral taxonomy level labelled in Ensembl as “vertebrates” or “jawed vertebrates”) to another gene itself orthologous to a human ISG. In total, our list includes 72 ancestral genes (Table I, Figure 4A).

**Figure 4.**
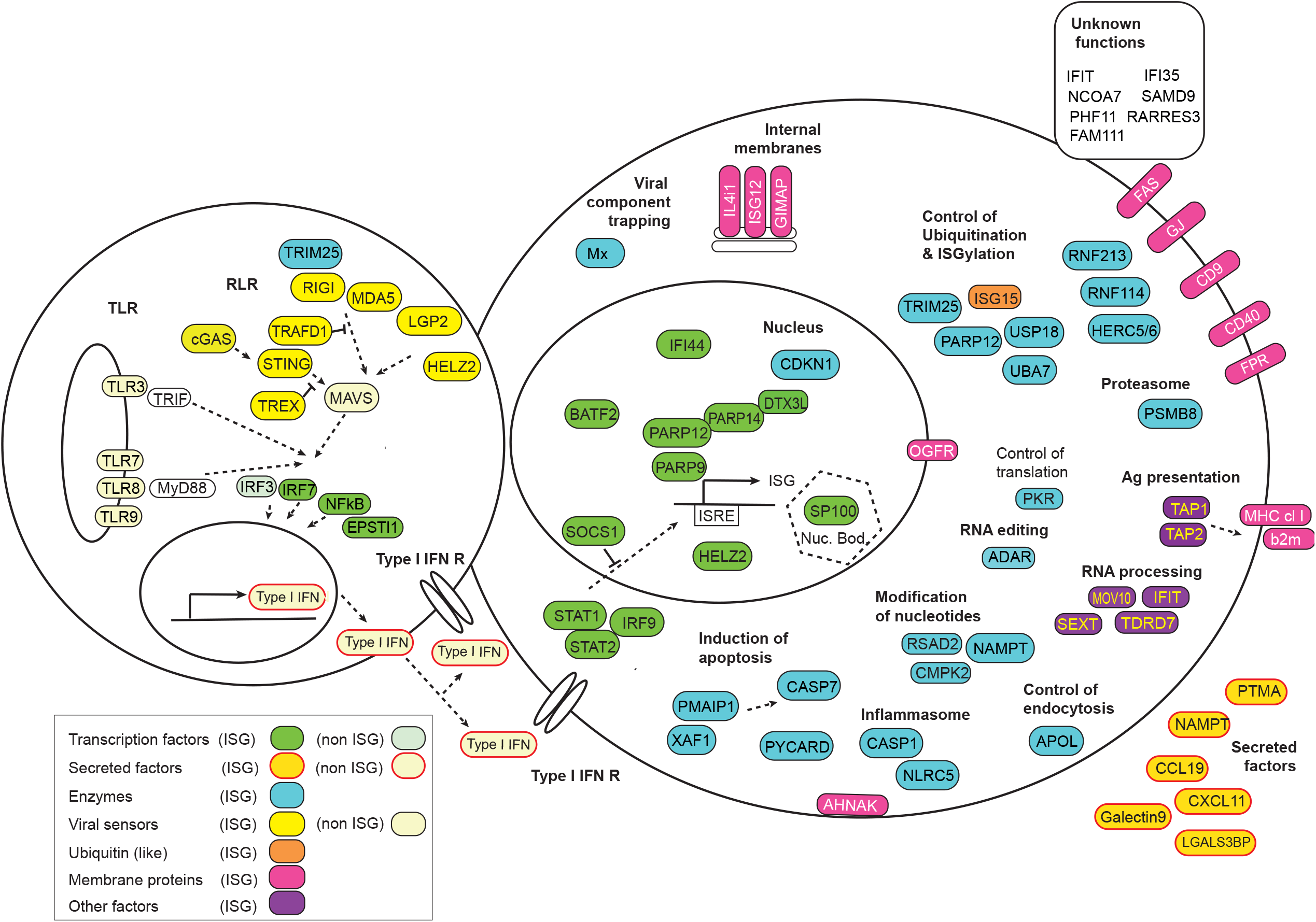
**Graphic overview of the ancestral ISG repertoire,** organized by functional modules.

Based on our orthology analysis, we propose new, more explicit names for many of the zebrafish ISGs with known human orthologues (in red on Table S5). This ancestral ISG core includes most ISGs with known functions. The IFN system of 450My ago seems fairly similar to the present one, particularly in its signaling components (Figure 4). Many ancestral genes have been duplicated independently in one or both lineages (Table S5), in addition to multiple ISGs apparently gained by either group (Figure 5).

**Figure 5.**
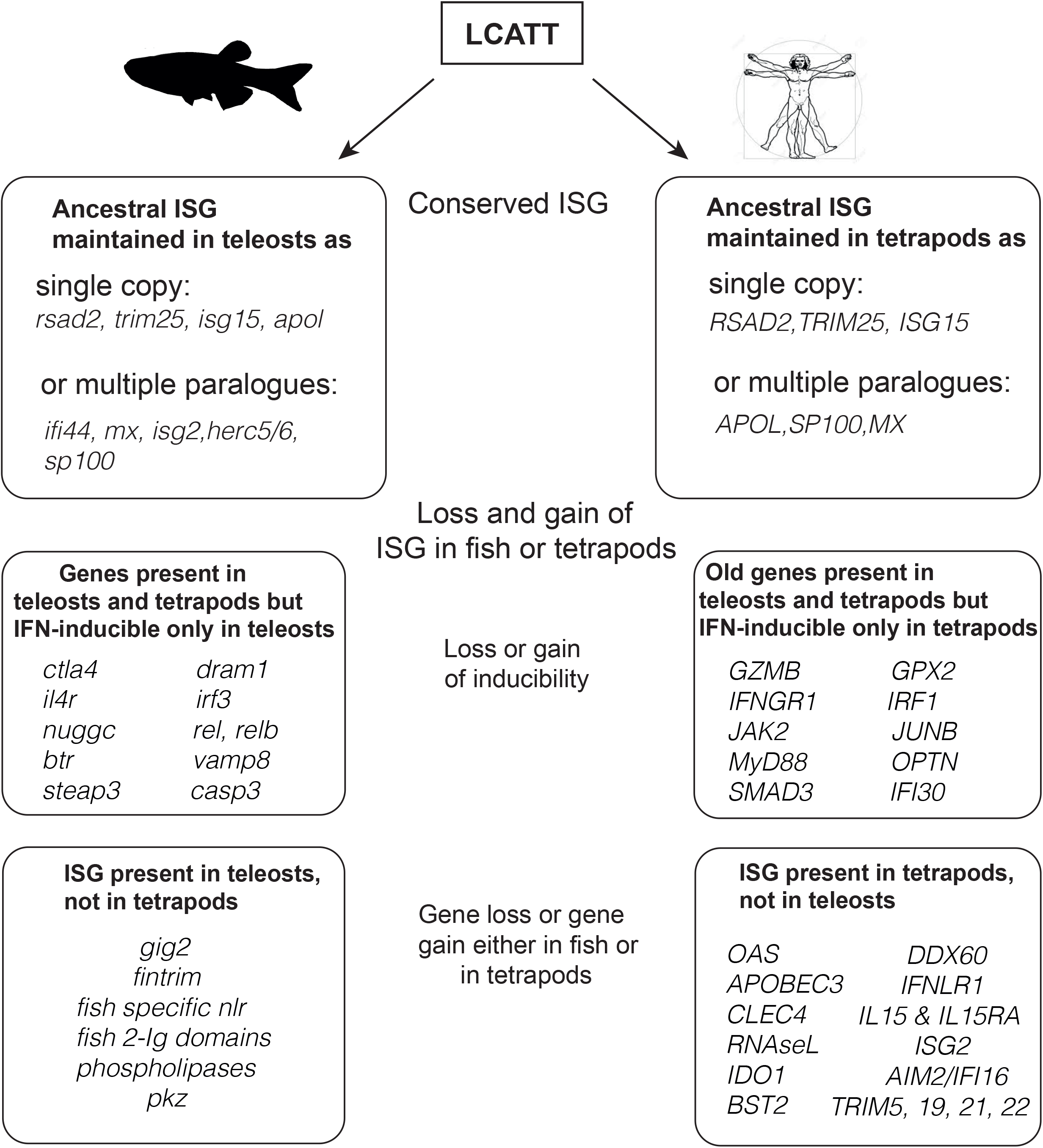
**Evolution of the ISG repertoire** since the LCATT (last common ancestor of Tetrapods and Teleosts). Genes names are given as examples, with no attempt to be exhaustive.

### IFN-downregulated genes

Two of the most strongly IFNφ1-downregulated genes (Table S1, bottom) were orthologous to human genes downregulated by type I IFNs, according to the Interferome database: plin1 (perilipin 1) and acox1 (palmytoil acyl-CoA oxidase 1). This suggests that downregulation of fatty acid oxidation pathway is an ancient feature of the IFN system.

Many IFNφ1 egulated genes were orthologous to a human gene in a 1-to-2 manner, with the two zebrafish paralogues having arisen during the teleost specific whole genome duplication (ohnologues). Systematically, only one of the two paralogues was downregulated.

Remarkably, no gene was downregulated by both IFNφ1 injection and CHIKV infection (Figure 3b, Table S3).

## Discussion

The zebrafish has become an important model to study host-pathogen interactions, particularly at its early life stages which are the most prone to live imaging and genetically tractable. Although its antiviral interferon genes and receptors are now well identified, knowledge of IFN-induced genes, or ISGs, was only partial. In this work, we used deep sequencing to characterize the transcriptomic response of the 3dpf zebrafish larva to recombinant IFNφ1, the first type I IFN identified in zebrafish and the most highly inducible one. We analyzed in parallel the response to an alphavirus inducing a strong type I IFN induction, ant the impact of IFN receptor knock-down on this response. From these different datasets, we established a comprehensive list of zebrafish ISGs. This list was compared to the human ISG repertoire, and a phylogenetic analysis was performed to approach the ancestral ISG repertoire of early vertebrates.

### 1. New insights and limitations of the work

A number of studies have identified genes induced by IFN or viral infections in fish (reviewed in (7)). However, very few global descriptions after treatment with recombinant type I IFN have been reported, using micro-arrays (19). Micro-array analyses are limited by probe choice, and are typically biased towards genes with known human homologues. RNAseq, by contrast, is mainly limited by the genome annotation quality and by the analysis method, and can be reanalyzed; this approach is thus more complete. Since the early zebrafish larva constitutes a reference model for investigating innate immune response, drug screening as well as for modelling diseases, we undertook a comprehensive description of the repertoire of ISG up-regulated at this developmental stage. Importantly, we previously reported a clear transcriptional response of zebrafish embryos to IFNφ1 as early as 24 hpf (Levraud et al. 2007); the responsiveness to type I IFNs is thus already well established at 3 dpf. We are aware that cells present in adult but not yet in larvae, notably those of the adaptive immune system such as lymphocytes and dendritic cells may express additional ISGs, which should be assessed in further work.

There are two groups of type I IFNs in teleost fish (40) with two different receptors (10). This study only addresses the ISG repertoire induced by IFNφ1 (a group 1 IFN) and it is possible that group 2 IFNs (IFNφ2 and IFNφ3) induce a different ISG subset. Determining this will require more studies; however, since CHIKV induces both IFNφ1 and IFNφ3 while *crfb1*&2 morpholinos target receptors for both type I IFN groups, IF φ therefore be found among CHIKV-induced, IFNR-dependent, but non IFNφ1 genes. Such genes (listed on Table S2, bottom) constituted about 30% of genes for which induction by CHIKV was impacted in morphants – and only about 15% if one also excludes genes for which induction by IFNφ1 is almost significant (Figure 3A and C). A previous report by López-Muñoz et al suggests differences in ISG induction, notably in kinetics, by different IFNφs (20).

### 2. Comparative and phylogenetic analysis of zebrafish ISGs

Our comparative and phylogenetic approach led to a tentative reconstruction of the ISG repertoire of the last common ancestor of teleosts and tetrapods (LCATT) that lived ∼450My ago and probably resembled the fossil osteichtyan Ligulalepis (41). To do so, we looked for human (co-)orthologue(s) of all zebrafish ISGs identified in our analysis. Based on available data compilations (30, 31), we then determined which one(s) of these human orthologues were themselves induced by type I IFN. In such cases, we considered that they most likely originated from an “ancestral” ISG, present in the LCATT. It is generally believed that the type I IFN system emerged during the early evolution of jawed vertebrates, since Chondrichtyans (rays, sharks and chimeras) but not Agnathans (lampreys and hagfish) possess typical type I IFN genes (4, 42). Hence, it is important to note that the IFN system had already evolved, expanded and standardized for more than 50My before our last common ancestor with zebrafish.

Approximately half of what we defined as ancestral ISGs are represented by 1-to-1 orthologues in zebrafish and human (Table S5, top rows) – a situation of practical interest, as the likelihood of conservation of gene function is highest in this case. These are either isolated genes (e.g. RSAD2, ISG15 or cGAS), or members of “old” families already stabilized in the LCATT (e.g., IRF7 and IRF9) (26). The situation is relatively similar for a few ancestral genes such as STAT1 or SOCS1, with one human orthologue and two zebrafish co-orthologues that arose during the teleost-specific whole genome duplication and were retained. In contrast, many other “young” families have clearly been subjected to further duplication during later evolution of fish or tetrapods, leading to orthology groups containing multiple ISG both in zebrafish and human, the most spectacular examples being the ISG12, IFIT and IFI44 families.

The frequency of orthology with a human gene is lower for ISGs (59%) than for the entire genome (71%). This is probably a consequence of the stronger evolutionary pressure of genes involved in the arms race with pathogens, as postulated by the Red Queen hypothesis (43). Similar mechanisms also explain the frequent and extensive gene duplications, as well as gene losses if some virus disappears, removing the corresponding selective pressure on a given ISG. Possibly, a greater diversity of aquatic viruses could further favor ISG retention and divergence after duplication, but few direct evidences are available.

In addition to the ancestral genes with true zebrafish and human orthologues, we added to this list a few genes with a more complex history, with a human and zebrafish ISGs that shared an ancestor at the basal vertebrate level (Table I and Table S5, bottom). These ancestral genes must have been duplicated in the LCATT genome; the teleost and tetrapod lineages then retained distinct paralogues. This comprises some genes whose evolutionary history is extremely difficult to trace due to multiple copies and extensive polymorphism, such as MHC class I genes. Here, only mhc1zba was found to be a zebrafish ISG, but this does not necessarily imply that other zebrafish MHC class I genes are not ISGs, as they may have been missed due to mapping issues; the strain we used (AB) is not the same as the one of the reference genome (Tü), and strain-specific divergences are considerable between strains for MHC class I, with deep evolutionary roots (44). Importantly, we did not define ancestral ISGs for zebrafish-human ISG pairs that appeared to be related at first glance, but, upon further analysis, were too distant; for example, zebrafish vamp5 and human VAMP8 are both ISGs, but share their last common ancestor at the Opistokontha level, before the split of fungi and animals, very long before the emergence of IFNs.

Nevertheless, the type I IFN system also includes very old genes that were already present in basal metazoans. The RNAseL/OAS module is a good example of such cases, being found across metazoans from mammals to sponges (45) -but lost in the fish branch. Another striking example is the cGAS-STING module recently identified in cnidaria (46). The implication of these genes in the antiviral immunity of basal branches of animals is unknown but certainly worth investigating. The main models for invertebrate immunity are flies and mosquitoes but they largely rely on RNAi mechanisms to contain viruses (3). Central signaling modules of the vertebrate IFN system such as TLR/NFκB and JAK/STAT, are also present in insects and in more distant metazoans, but they induce different set of genes with other functions (47).

Additionally, a few important genes do not meet our criteria for “ancestral” ISG because they are not typically inducible either in zebrafish or in human (Figure 5). For example, irf3 is an ISG in fish but not in human, while it is the reverse for JAK2. Hence, our list of ancestral ISG is likely not complete, but it provides a core repertoire pointing to most fundamental factors of the vertebrate innate antiviral arsenal.

A relatively large number of ISGs have no orthologue in the other lineage, such as human APOBEC3, RNASEL, OAS, AIM2 (Figure 5). Similarly, many fish-specific ISG likely have been co-opted by the IFN pathway during fish evolution. In this case, they do not have clear orthologues in human and other tetrapods (as for finTRIMs and nlr-B30.2), or their orthologue(s) have no link with the type I IFN system. The finTRIM family contains the largest number of zebrafish ISGs of any family, ancestral or not. Interestingly, ISGs are found only among the recently diversified, species-specific finTRIMs; the most basal members (ftr82-84), well-conserved among fish, were not found here to be induced by IFNφ1 or by CHIKV, consistent with previous studies (48). Nevertheless, ftr83 appears to mediate protection especially in the gill region by stimulating local *ifnphi1*expression (49). The diversity and evolution under positive selection of the IFN-inducible finTRIMs evoke viral recognition (22), yet their functions remain unclear.

The co-optation of new genes in the ISG repertoire may be operated quickly and in a group-specific manner, by introduction of sequence motifs in the regulatory sequences, for example via retroviral insertion (50, 51). However, we cannot exclude that these branch-specific ISGs are in fact ancestral, but lost in one of the two lineages; this is the case for Gig2 genes, which are present in the Coelacanth genome as well as in fish, and thus were lost in tetrapods. Thus, our repertoire of ancestral ISGs is underestimated because we cannot include the lineage-specific losses.

Do ancestral ISGs identified here define a minimal but complete set of response elements from recognition to elimination of invading viruses? Probably not, as this ancestral core group of ISGs was backed up by more ISGs is any species, including the LCATT. For example, the absence of the well-known OAS/RNAseL module genes in fish (and therefore in our list of conserved ancestral ISGs) is puzzling, and one could predict that other fish genes have taken over similar functions. Similarly, APOBEC3 genes are absent in fish, and maybe their RNA-editing mechanisms are mediated by other genes, possibly by ADAR1.

Ancestral ISGs encode very diverse proteins in localization and function (Figure 4). We provide an extended discussion of their classification below.

### 4. Characterization of the IFN independent response to CHIKV infection

Knowing the repertoire of ISG also offers the possibility to identify genes that are induced by viral infection independently of the type I IFN pathway. While a subset of ISGs can be induced via IFN-dependent and -independent pathways in human and fish - for example rsad2 (12). Thus, IRF3 dependent, type I IFN independent induction of many ISG by particular viruses has been described (52).

However, about half of the genes upregulated by CHIKV were not induced by IFNϕ1 injection, and most were not affected by IFNR knockdown. Notably, three gene sets stand out in this list: (1) components of the complement cascade, that are known to play a role in antiviral defense; (2) cytokines including some CC and CXC chemokines as well as the type I IFN themselves, which do not appear to be strongly auto/cross inducible and (3) many btr and ftr TRIM E3 ligases as well as multiple proteasome components. Interestingly too, irf1b, the zebrafish orthologue of IRF1 (a human ISG), is CHIKV-inducible, but not in a IFNR-dependent manner – consistent with previous work (26) - and was not induced by IFNϕ1 Many other genes of unknown function also share the same induction pattern, and would certainly be worth investigating. A strong redundancy of antiviral pathways has certainly been selected during evolution, since viruses have developed multiple strategies of immune subversion.

Contrary to what was observed with upregulated genes, there was no overlap between gene sets downregulated by IFNϕ1 and by CHIKV. This remarkable difference could be due to the alternative inflammatory response induced by the virus besides type I IFNs, or to kinetic differences.

### 5. Classification of ancestral ISGs

The ancestral ISG presented in Table I can be classified based on molecular functions: sensors, transcription factors and other signal transduction factors, secreted factors, enzymes including ubiquitination factors, and membrane receptors, which we discuss below. The antiviral mechanisms described in human or in other mammalian systems also provide hints about the likely conserved mode of action of these factors.

#### Transcription factors

Many members of the list appear to have DNA binding capacity and may be classified as transcription factors.

* Three IRF (3, 7, 9) and two STAT(1, 2) constitute fundamental components of the type I pathway signaling, and were already present in the LCA of fish and mammals.
* BATF2 is a member of the AP-1/ATF family transcription factors that controls the differentiation of immune cells and play key regulatory roles in immune responses. BATF2 promotes TLR7-induced Th1 responses (53).
* SP100 is a tumor suppressor and a major constituent of the PML bodies controlling transcription and/or chromatin conformation.
* HELZ2 is a helicase that acts as a transcriptional coactivator for a number of nuclear receptors, including AHR, a nuclear receptor regulating lipid metabolism and the susceptibility to dengue virus (54).
* transcription coactivators. PARPs can act as transcriptional co-activators and potentiate induction of many ISG (55). The multiple *ifi44* zebrafish genes counts 7 ISG among 19 members, but their two human co-orthologues are induced by type I IFN. Located in the nucleus, IFI44 binds and blocks the HIV1 LTR promoter (56). However, the numerous zebrafish *ifi44* probably have sub-functionalized, and mediate multiple antiviral mechanisms.

#### Sensors and related genes

The helicases RIG-I, LPGP2 and IFIH1 (aka MDA5) stand as primary ISGs encoding viral sensors. Besides, as a cytoplasmic helicase HELZ2 might also play a sensor role. In keeping with this, TREX proteins have a 3’-to-5’ DNA exonuclease activity that is important to block the sting-dependent initiation of IFN responses by DNA fragments from endogenous retroviruses and elements (57).

#### Enzymes

Besides transcription factors, enzymes are the most important category of ancestral ISG. They may play a role in signaling or have a direct antiviral activity.

* Poly-ADP-ribose polymerase (PARP) are involved in many cellular processes, from regulation of chromatin conformation to transcription control, and several PARP also are induced by infection and inflammation. PARP are represented by parp9, parp12 and parp14 among ancestral ISG. Strikingly, these three PARP are part of a nuclear complex, with the E3 ubiquitin ligase encoded by dtx3l that is also an ancestral ISG that promoting RNA Pol II recruitment at IRF3-dependent promoters (55). Our data showing that key components of this complex are part of the essential type I IFN system underscore its importance in the core antiviral response. Besides, other activities of PARP may be involved in antiviral mechanisms; for example PARP12 mediates ADP-ribosylation of Zika virus NS1 and NS3, leading to their degradation by the proteasome (58). The ADP-ribo-hydrolase encoded by the Chikungunya virus, that is required for its virulence, is another hint of the central importance of these enzymes in antiviral defense (59).
* Several E3 Ubiquitin ligases were found among ancestral ISG, including trim25, usp18, rnf114, dtx3l. The mechanisms through which they exert antiviral activity or regulate the response are not fully resolved. The critical role of trim25 in RIGI activation, and its capacity of ISGylation (60) have been well documented in fish and mammals. isg15, an ubiquitin like protein, is also an ancestral ISG playing a central role in the type I IFN pathway in fish and mammals (7, 60) via multiple mechanisms.
* The pro-apoptotic caspase casp7 possess type I IFN induced orthologues in zebrafish and human. Interestingly, ancestral ISG also comprise pmaip1 that promotes caspase activation and apoptosis via modifications of the mitochondrial membrane, and xiaf1, a negative regulator of members of inhibitor of apoptosis proteins. Taken together, these observations indicate that the ancestral type I IFN system comprised a pro-apoptotic module.
* The pro-inflammatory caspase casp1, is also an ancestral ISG, as is pycard which encodes ASC, the major scaffold protein of the canonical inflammasome. Induction of the inflammasome is thus an ancestral property of the IFN response. Many upstream sensors of the inflammasome are IFN-inducible, but they are generally divergent in the two lineages, nlrc5 being the only ancestral ISG.
* Rsad2 (aka viperin) is an enzyme with a direct antiviral function, that catalyzes the conversion of CTP to a completely new ribonucleoside, the 3’-deoxy-3’,4’-didehydro-CTP (ddhCTP) acting as a terminator of RNA synthesis (61). Interestingly, both ancestral ISG rsad2 and the nucleotide modifier cpmk2 are located very close to each other in the genome in fish as well as in mammals; likely forming a conserved functional antiviral unit.
* Adenosine deaminases acting on double-stranded RNA (ADARs) deaminate adenosine to produce inosine in double-stranded RNA structures, regulating the inflammation induced by such molecules (62). Accordingly, loss of function of adar in zebrafish larvae leads to brain inflammation in a model of Aicardi-Goutières syndrome, suggesting a key regulatory role of this gene during type I IFN response (63).
* Protein Kinase R (PKR, encoded by *eif2ak2*) is activated by dsRNA (and thus could have been listed above as a sensor), leading phosphorylation of EIF2α, and to inhibition of protein synthesis and viral replication. Many viruses encode PKR inhibitors of this cornerstone antiviral factor that also affects transcription factors like IRF1, STATs, and NF-kappaB and upregulates many genes including β2microglobulin and isg15 (64). Interestingly, the other ancestral ISG epsti1 can activate PKR promoters and induce PKR-dependent genes in human (65), questioning whether pkr and epsti1 may have been functionally coupled from the LCATT. Fish possess a lineage specific paralogue of PKR called PKZ, which detects Z-DNA (66).

#### Secreted factors

* In humans and mice, ccl19 is implicated in lymphocyte migration and is important to define compartments within lymphoid tissues. In rainbow trout, one of the six ccl19 paralogues present in the genome participate to antiviral immunity though promotion of mucosal and central CD8+ T cell response (67).
* Some of the fish homologues of murine and human IFN inducible CXC chemokines – ie, CXCL9-11, which bind CXCR3, a receptor expressed by various leukocyte, including some T cells, macrophages, and dendritic cell subsets – are also up-regulated by IFNφ in zebrafish larvae. These genes have been largely expanded in fish, and two lineages of CXCL11 have been recently distinguished, both closely related to the mammalian CXCL9-11 (68, 69). The up-regulated cxcl11.3 (aka cxc66 or cxcl11ac) identified in this work belongs to the lineage 1. The zebrafish has three cxcr3 paralogues, and receptor-ligand binding, tested for three other zebrafish cxcl11 ligands, does not follow ligand lineage (70), so the receptor(s) of this ISG remains to be identified experimentally.
* Another soluble factor up-regulated by type I IFN and viral infection in fish is galectin9 (this work, and (13)). In mammals, Galectin9 is involved in multiple mechanisms of antiviral immunity. For example, it is a potent factor against HCMV because it blocks the entry of the virus in target cells (71). Galectin9 can also regulates HIV transcription, and induces the expression of the deaminase APOBEC3G, a potent antiviral factor (72). Besides, the galectin-9 receptor TIM3 is implicated in the control of Th1 cells (73).

#### Membrane proteins

* While zebrafish and human mhc class I are not direct orthologues, mhc class I genes are in the list, with β2microglobulin and the peptide transporters tap-1 and tap-2, as well as homologues of TAPBP and proteasome subunits, indicating that this pathway is a fundamental component of the type I IFN system.
* Other important membrane proteins in the list are tetraspanins of the CD9 family, that regulate degranulation of myeloid subsets and secretion of cytokines, hence constitute key players in inflammation (74).
* Zebrafish possess eight isg12 genes located in tandem, of which six were highly inducible by IFNϕ1 and by CHIKV. Their human ISG orthologues IFI6 and IFI27 (aka ISG12A) are internal membrane proteins stabilizing ER membrane and preventing the formation of flavivirus-induced ER membrane invaginations (75) or destabilize mitochondrial membrane and promote apoptosis (76). In fact, IFI27 can also recruit a E3 ubiquitin ligase and targets HCV NS5 protein to degradation (77), illustrating the potential diversity of antiviral mechanisms mediated by members of this family.
*APOL1 affects endocytosis and promotes an expansion of the lysosomal compartment, favoring for example the degradation of the HIV-1 protein Vif (78).

#### ISG with unknown functions or unknown antiviral mechanisms

Even in human and mice, the basis of antiviral activity of certain ISGs remains completely unknown. For example, the effects of PHF11, RNF114, or SAMD9 are elusive. In the latter, a DNA/RNA-binding AlbA, a NTPase, and a OB domain with predicted RNA-binding properties suggest a link with nucleic acid metabolism or sensing (79). These very old ISG with counterparts found across Metazoa and even in procaryotes, are key restriction factors of poxviruses (80).

## 6. Conclusions

Antiviral genes are well known to evolve very fast, as postulated by the Red Queen hypothesis, under strong pressure from pathogens. This is indeed illustrated by the large number of ISGs that are either fish- or mammal-specific. Nevertheless, our data define a surprisingly stable set of core ISGs, that were apparently co-opted into the new IFN system of early vertebrates about 500My ago, and have been maintained for the last 450My both in fish and tetrapods. The full list of zebrafish ISG provides a powerful reference to characterize the subtle interactions between viruses and the host response, including redundancy of immune pathways and viral subversion mechanisms. It also constitutes a valuable resource for the study of autoinflammatory disease using the emerging zebrafish model.

## Acknowledgements

We thank Jean-Yves Coppee and Caroline Proux from the transcriptomics platform of Institut Pasteur, for generation and sequencing of RNA libraries. We are indebted to Rune Hartmann (Aarhus, Denmark) for recombinant zebrafish interferon and stimulating discussion. We thank Emma Colucci and Pedro Hernandez-Cerda for critical reading of the manuscript.

## References

1. Schoggins, J. W., and C. M. Rice. 2011. Interferon-stimulated genes and their antiviral effector functions. Curr. Opin. Virol. 1: 519–525.

2. Backes, S., R. A. Langlois, S. Schmid, A. Varble, J. V. Shim, D. Sachs, and B. R. TenOever. 2014. The Mammalian Response to Virus Infection Is Independent of Small RNA Silencing. Cell Rep. 8: 114–125.

3. Guo, Z., Y. Li, and S. W. Ding. 2019. Small RNA-based antimicrobial immunity. Nat. Rev. Immunol. 19: 31–44.

4. Secombes, C. J., and J. Zou. 2017. Evolution of interferons and interferon receptors. Front. Immunol. 8: 209.

5. Redmond, A. K., J. Zou, C. J. Secombes, D. J. Macqueen, and H. Dooley. 2019. Discovery of All Three Types in Cartilaginous Fishes Enables Phylogenetic Resolution of the Origins and Evolution of Interferons. Front. Immunol. 10: 1558.

6. Zou, J., and C. J. Secombes. 2011. Teleost fish interferons and their role in immunity. Dev. Comp. Immunol. 35: 1376–1387.

7. Langevin, C., E. Aleksejeva, G. Passoni, N. Palha, J. P. Levraud, and P. Boudinot. 2013. The antiviral innate immune response in fish: Evolution and conservation of the IFN system. J. Mol. Biol. 425: 4904–4920.

8. Boudinot, P., C. Langevin, C. J. Secombes, and J.-P. Levraud. 2016. The peculiar characteristics of fish type I interferons. Viruses 8.

9. Robertsen, B. 2018. The role of type I interferons in innate and adaptive immunity against viruses in Atlantic salmon. Dev. Comp. Immunol. 80: 41–52.

10. Aggad, D., M. Mazel, P. Boudinot, K. E. Mogensen, O. J. Hamming, R. Hartmann, S. Kotenko, P. Herbomel, G. Lutfalla, and J.-P. Levraud. 2009. The two groups of zebrafish virus-induced interferons signal via distinct receptors with specific and shared chains. J. Immunol. 183: 3924–3931.

11. Staeheli, P., Y. X. Yu, R. Grob, and O. Haller. 1989. A double-stranded RNA-inducible fish gene homologous to the murine influenza virus resistance gene Mx. Mol. Cell. Biol. 9: 3117–21.

12. Boudinot, P., P. Massin, M. Blanco, S. Riffault, and A. Benmansour. 1999. vig-1, a new fish gene induced by the rhabdovirus glycoprotein, has a virus-induced homologue in humans and shares conserved motifs with the MoaA family. J. Virol. 73: 1846–1852.

13. O’Farrell, C., N. Vaghefi, and M. Cantonnet. 2002. Survey of Transcript Expression in Rainbow Trout Leukocytes Reveals a Major Contribution of Interferon-Responsive Genes in the Early Response to a Rhabdovirus Infection. J. Virol. 76: 8040–8049.

14. Levraud, J.-P., P. Boudinot, I. Colin, A. Benmansour, N. Peyrieras, P. Herbomel, and G. Lutfalla. 2007. Identification of the zebrafish IFN receptor: implications for the origin of the vertebrate IFN system. J. Immunol. 178: 4385–4394.

15. Lutfalla, G., H. R. Crollius, N. Stange-Thomann, O. Jaillon, K. Mogensen, and D. Monneron. 2003. Comparative genomic analysis reveals independent expansion of a lineage-specific gene family in vertebrates: The class II cytokine receptors and their ligands in mammals and fish. BMC Genomics 4: 29.

16. Altmann, S. M., M. T. Mellon, D. L. Distel, and C. H. Kim. 2003. Molecular and Functional Analysis of an Interferon Gene from the Zebrafish, Danio rerio. J. Virol. 77: 1992–2002.

17. Robertsen, B., V. Bergan, T. Røkenes, R. Larsen, and A. Albuquerque. 2003. Atlantic Salmon Interferon Genes: Cloning, Sequence Analysis, Expression, and Biological Activity. J. Interf. Cytokine Res. 23: 601–612.

18. Purcell, M. K., K. M. Nichols, J. R. Winton, G. Kurath, G. H. Thorgaard, P. Wheeler, J. D. Hansen, R. P. Herwig, and L. K. Park. 2006. Comprehensive gene expression profiling following DNA vaccination of rainbow trout against infectious hematopoietic necrosis virus. Mol. Immunol. 43: 2089–2106.

19. Martin, S. A. M., J. B. Taggart, P. Seear, J. E. Bron, R. Talbot, A. J. Teale, G. E. Sweeney, B. Høyheim, D. F. Houlihan, D. R. Tocher, J. Zou, and C. J. Secombes. 2007. Interferon type I and type II responses in an Atlantic salmon (Salmo salar) SHK-1 cell line by the salmon TRAITS/SGP microarray. Physiol. Genomics 32: 33–44.

20. López-Muñoz, A., F. J. Roca, J. Meseguer, and V. Mulero. 2009. New insights into the evolution of IFNs: zebrafish group II IFNs induce a rapid and transient expression of IFN-dependent genes and display powerful antiviral activities. J. Immunol. 182: 3440–3449.

21. Rothenburg, S., N. Deigendesch, M. Dey, T. E. Dever, and L. Tazi. 2008. Double-stranded RNA-activated protein kinase PKR of fishes and amphibians: varying the number of double-stranded RNA binding domains and lineage-specific duplications. BMC Biol. 6: 12.

22. van der Aa, L. M., J.-P. Levraud, M. Yahmi, E. Lauret, V. Briolat, P. Herbomel, A. Benmansour, and P. Boudinot. 2009. A large new subset of TRIM genes highly diversified by duplication and positive selection in teleost fish. BMC Biol. 7: 7.

23. Zhang, Y. B., and J. F. Gui. 2004. Identification and expression analysis of two IFN-inducible genes in crucian carp (Carassius auratus L.). Gene 60: 1–9.

24. Zhang, Y. B., T. K. Liu, J. Jiang, J. Shi, Y. Liu, S. Li, and J. F. Gui. 2013. Identification of a Novel Gig2 Gene Family Specific to Non-Amniote Vertebrates. PLoS One 8: e60588.

25. Boudinot, P., S. Salhi, M. Blanco, and A. Benmansour. 2001. Viral haemorrhagic septicaemia virus induces vig −2, a new interferon-responsive gene in rainbow trout. Fish Shellfish Immunol. 11: 383–397.

26. Briolat, V., L. Jouneau, R. Carvalho, N. Palha, C. Langevin, P. Herbomel, O. Schwartz, H. P. Spaink, J.-P. Levraud, and P. Boudinot. 2014. Contrasted Innate Responses to Two Viruses in Zebrafish: Insights into the Ancestral Repertoire of Vertebrate IFN-Stimulated Genes. J. Immunol. 192: 4328–41.

27. Ludwig, M., N. Palha, C. Torhy, V. Briolat, E. Colucci-Guyon, M. Brémont, P. Herbomel, P. Boudinot, and J. P. Levraud. 2011. Whole-body analysis of a viral infection: Vascular endothelium is a primary target of Infectious Hematopoietic Necrosis Virus in zebrafish larvae. PLoS Pathog. 7: e1001269.

28. Palha, N., F. Guivel-Benhassine, V. Briolat, G. Lutfalla, M. Sourisseau, F. Ellett, C. H. Wang, G. J. Lieschke, P. Herbomel, O. Schwartz, and J. P. Levraud. 2013. Real-Time Whole-Body Visualization of Chikungunya Virus Infection and Host Interferon Response in Zebrafish. PLoS Pathog. 9: e1003619.

29. Howe, K., M. D. Clark, C. F. Torroja, J. Torrance, C. Berthelot, M. Muffato, J. E. Collins, S. Humphray, K. McLaren, L. Matthews, S. McLaren, I. Sealy, M. Caccamo, C. Churcher, C. Scott, J. C. Barrett, R. Koch, G.-J. Rauch, S. White, W. Chow, B. Kilian, L. T. Quintais, J. a Guerra-Assunção, Y. Zhou, Y. Gu, J. Yen, J.-H. Vogel, T. Eyre, S. Redmond, R. Banerjee, J. Chi, B. Fu, E. Langley, S. F. Maguire, G. K. Laird, D. Lloyd, E. Kenyon, S. Donaldson, H. Sehra, J. Almeida-King, J. Loveland, S. Trevanion, M. Jones, M. Quail, D. Willey, A. Hunt, J. Burton, S. Sims, K. McLay, B. Plumb, J. Davis, C. Clee, K. Oliver, R. Clark, C. Riddle, D. Elliot, D. Eliott, G. Threadgold, G. Harden, D. Ware, S. Begum, B. Mortimore, B. Mortimer, G. Kerry, P. Heath, B. Phillimore, A. Tracey, N. Corby, M. Dunn, C. Johnson, J. Wood, S. Clark, S. Pelan, G. Griffiths, M. Smith, R. Glithero, P. Howden, N. Barker, C. Lloyd, C. Stevens, J. Harley, K. Holt, G. Panagiotidis, J. Lovell, H. Beasley, C. Henderson, D. Gordon, K. Auger, D. Wright, J. Collins, C. Raisen, L. Dyer, K. Leung, L. Robertson, K. Ambridge, D. Leongamornlert, S. McGuire, R. Gilderthorp, C. Griffiths, D. Manthravadi, S. Nichol, G. Barker, S. Whitehead, M. Kay, J. Brown, C. Murnane, E. Gray, M. Humphries, N. Sycamore, D. Barker, D. Saunders, J. Wallis, A. Babbage, S. Hammond, M. Mashreghi-Mohammadi, L. Barr, S. Martin, P. Wray, A. Ellington, N. Matthews, M. Ellwood, R. Woodmansey, G. Clark, J. D. Cooper, J. Cooper, A. Tromans, D. Grafham, C. Skuce, R. Pandian, R. Andrews, E. Harrison, A. Kimberley, J. Garnett, N. Fosker, R. Hall, P. Garner, D. Kelly, C. Bird, S. Palmer, I. Gehring, A. Berger, C. M. Dooley, Z. Ersan-Ürün, C. Eser, H. Geiger, M. Geisler, L. Karotki, A. Kirn, J. Konantz, M. Konantz, M. Oberländer, S. Rudolph-Geiger, M. Teucke, C. Lanz, G. Raddatz, K. Osoegawa, B. Zhu, A. Rapp, S. Widaa, C. Langford, F. Yang, S. C. Schuster, N. P. Carter, J. Harrow, Z. Ning, J. Herrero, S. M. J. Searle, A. Enright, R. Geisler, R. H. a Plasterk, C. Lee, M. Westerfield, P. J. de Jong, L. I. Zon, J. H. Postlethwait, C. Nüsslein-Volhard, T. J. P. Hubbard, H. Roest Crollius, J. Rogers, and D. L. Stemple. 2013. The zebrafish reference genome sequence and its relationship to the human genome. Nature 496: 498–503.

30. Schoggins, J. W., S. J. Wilson, M. Panis, M. Y. Murphy, C. T. Jones, P. Bieniasz, and C. M. Rice. 2011. A diverse range of gene products are effectors of the type I interferon antiviral response. Nature 472: 481–5.

31. Rusinova, I., S. Forster, S. Yu, A. Kannan, M. Masse, H. Cumming, R. Chapman, and P. J. Hertzog. 2013. INTERFEROME v2.0: An updated database of annotated interferon-regulated genes. Nucleic Acids Res. 41: 1040–1046.

32. Levraud, J.-P., E. Colucci-Guyon, M. J. Redd, G. Lutfalla, and P. Herbomel. 2008. In vivo analysis of zebrafish innate immunity. Methods Mol. Biol. 415: 337–363.

33. Biacchesi, S., M. LeBerre, A. Lamoureux, Y. Louise, E. Lauret, P. Boudinot, and M. Brémont. 2009. Mitochondrial antiviral signaling protein plays a major role in induction of the fish innate immune response against RNA and DNA viruses. J. Virol. 83: 7815–27.

34. Letunic, I., and P. Bork. 2018. 20 years of the SMART protein domain annotation resource. Nucleic Acids Res. 46: D493–496.

35. Boudinot, P., L. M. van der Aa, L. Jouneau, L. Pasquier, P. Pontarotti, V. Briolat, A. Benmansour, and J. P. Levraud. 2011. Origin and evolution of TRIM proteins: New insights from the complete TRIM repertoire of zebrafish and pufferfish. PLoS One 6: e22022.

36. Howe, K., P. H. Schiffer, J. Zielinski, T. Wiehe, G. K. Laird, J. C. Marioni, O. Soylemez, F. Kondrashov, and M. Leptin. 2016. Structure and evolutionary history of a large family of NLR proteins in the zebrafish. Open Biol. 6: 160009.

37. Stein, C., M. Caccamo, G. Laird, and M. Leptin. 2007. Conservation and divergence of gene families encoding components of innate immune response systems in zebrafish. Genome Biol. 8: R251.

38. Laing, K. J., M. K. Purcell, J. R. Winton, and J. D. Hansen. 2008. A genomic view of the NOD-like receptor family in teleost fish: Identification of a novel NLR subfamily in zebrafish. BMC Evol. Biol. 8: 42.

39. Zhang, F., Y. Tao, Z. Zhang, X. Guo, P. An, Y. Shen, Q. Wu, Y. Yu, and F. Wang. 2012. Metalloreductase steap3 coordinates the regulation of iron homeostasis and inflammatory responses. Haematologica 97: 1826–1835.

40. Zou, J., C. Tafalla, J. Truckle, and C. J. Secombes. 2007. Identification of a second group of type I IFNs in fish sheds light on IFN evolution in vertebrates. J. Immunol. 179: 3859–3871.

41. Clement, A. M., B. King, S. Giles, B. Choo, P. E. Ahlberg, G. C. Young, and J. A. Long. 2018. Neurocranial anatomy of an enigmatic Early Devonian fish sheds light on early osteichthyan evolution. Elife 7: e34349.

42. Venkatesh, B., A. P. Lee, V. Ravi, A. K. Maurya, M. M. Lian, J. B. Swann, Y. Ohta, M. F. Flajnik, Y. Sutoh, M. Kasahara, S. Hoon, V. Gangu, S. W. Roy, M. Irimia, V. Korzh, I. Kondrychyn, Z. W. Lim, B. H. Tay, S. Tohari, K. W. Kong, S. Ho, B. Lorente-Galdos, J. Quilez, T. Marques-Bonet, B. J. Raney, P. W. Ingham, A. Tay, L. W. Hillier, P. Minx, T. Boehm, R. K. Wilson, S. Brenner, and W. C. Warren. 2014. Elephant shark genome provides unique insights into gnathostome evolution. Nature 505: 174–179.

43. tenOever, B. R. 2016. The Evolution of Antiviral Defense Systems. Cell Host Microbe 19: 142–149.

44. McConnell, S. C., K. M. Hernandez, D. J. Wcisel, R. N. Kettleborough, D. L. Stemple, J. A. Yoder, J. Andrade, and J. L. O. de Jong. 2016. Alternative haplotypes of antigen processing genes in zebrafish diverged early in vertebrate evolution. Proc. Natl. Acad. Sci. 113: E5014–E5023.

45. Päri, M., A. Kuusksalu, A. Lopp, T. Reintamm, J. Justesen, and M. Kelve. 2007. Expression and characterization of recombinant 2,5 - oligoadenylate synthetase from the marine sponge Geodia cydonium. FEBS J. 274: 3462–3474.

46. Kranzusch, P. J., S. C. Wilson, A. S. Y. Lee, J. M. Berger, J. A. Doudna, and R. E. Vance. 2015. Ancient Origin of cGAS-STING Reveals Mechanism of Universal 2’,3’ cGAMP Signaling. Mol. Cell 59: 891–903.

47. Liongue, C., R. Sertori, and A. C. Ward. 2016. Evolution of Cytokine Receptor Signaling. J. Immunol. 197: 11–18.

48. Langevin, C., J. P. Levraud, and P. Boudinot. 2019. Fish antiviral tripartite motif (TRIM) proteins. Fish Shellfish Immunol. 86: 724–733.

49. Langevin, C., E. Aleksejeva, A. Houel, V. Briolat, C. Torhy, A. Lunazzi, J.-P. Levraud, and P. Boudinot. 2017. FTR83, a member of the large fish-specific finTRIM family, triggers IFN pathway and counters viral infection. Front. Immunol. 8: 617.

50. Chuong, E. B., N. C. Elde, and C. Feschotte. 2016. Regulatory evolution of innate immunity through co-option of endogenous retroviruses. Science (80-.). 351: 1083–7.

51. Arnold, M. L., and K. Kunte. 2017. Adaptive Genetic Exchange: A Tangled History of Admixture and Evolutionary Innovation. Trends Ecol. Evol. 32: 601–611.

52. Ashley, C. L., A. Abendroth, B. P. McSharry, and B. Slobedman. 2019. Interferon-Independent Upregulation of Interferon-Stimulated Genes during Human Cytomegalovirus Infection is Dependent on IRF3 Expression. Viruses 11: E246.

53. Kanemaru, H., F. Yamane, K. Fukushima, T. Matsuki, T. Kawasaki, I. Ebina, K. Kuniyoshi, H. Tanaka, K. Maruyama, K. Maeda, T. Satoh, and S. Akira. 2017. Antitumor effect of Batf2 through IL-12 p40 up-regulation in tumor-associated macrophages. Proc. Natl. Acad. Sci. 114: E7331–E7340.

54. Fusco, D. N., H. Pratt, S. Kandilas, S. S. Y. Cheon, W. Lin, D. A. Cronkite, M. Basavappa, K. L. Jeffrey, A. Anselmo, R. Sadreyev, C. Yapp, X. Shi, J. F. O’Sullivan, R. E. Gerszten, T. Tomaru, S. Yoshino, T. Satoh, and R. T. Chung. 2017. HELZ2 is an IFN effector mediating suppression of dengue virus. Front. Microbiol. 8: 240.

55. Caprara, G., E. Prosperini, V. Piccolo, G. Sigismondo, A. Melacarne, A. Cuomo, M. Boothby, M. Rescigno, T. Bonaldi, and G. Natoli. 2018. PARP14 Controls the Nuclear Accumulation of a Subset of Type I IFN–Inducible Proteins. J. Immunol. 200: 2439–2454.

56. Power, D., N. Santoso, M. Dieringer, J. Yu, H. Huang, S. Simpson, I. Seth, H. Miao, and J. Zhu. 2015. IFI44 suppresses HIV-1 LTR promoter activity and facilitates its latency. Virology 481: 142–50.

57. Yan, N. 2017. Immune Diseases Associated with TREX1 and STING Dysfunction. J. Interf. Cytokine Res. 37: 198–206.

58. Li, L., H. Zhao, P. Liu, C. Li, N. Quanquin, X. Ji, N. Sun, P. Du, C. F. Qin, N. Lu, and G. Cheng. 2018. PARP12 suppresses Zika virus infection through PARP-dependent degradation of NS1 and NS3 viral proteins. Sci. Signal. 11: eaas9332.

59. McPherson, R. L., R. Abraham, E. Sreekumar, S.-E. Ong, S.-J. Cheng, V. K. Baxter, H. A. V. Kistemaker, D. V. Filippov, D. E. Griffin, and A. K. L. Leung. 2017. ADP-ribosylhydrolase activity of Chikungunya virus macrodomain is critical for virus replication and virulence. Proc. Natl. Acad. Sci. 114: 16666–1671.

60. Langevin, C., L. M. van der Aa, a Houel, C. Torhy, V. Briolat, a Lunazzi, a Harmache, M. Bremont, J.-P. Levraud, and P. Boudinot. 2013. Zebrafish ISG15 exerts a strong antiviral activity against RNA and DNA viruses and regulates the interferon response. J. Virol. 87: 10025–36.

61. Gizzi, A. S., T. L. Grove, J. J. Arnold, J. Jose, R. K. Jangra, S. J. Garforth, Q. Du, S. M. Cahill, N. G. Dulyaninova, J. D. Love, K. Chandran, A. R. Bresnick, C. E. Cameron, and S. C. Almo. 2018. A naturally occurring antiviral ribonucleotide encoded by the human genome. Nature 558: 610–614.

62. Samuel, C. E. 2019. Adenosine deaminase acting on RNA (ADAR1), a suppressor of double-stranded RNA-triggered innate immune responses. J. Biol. Chem. 294: 1710–1720.

63. Kasher, P. R., E. M. Jenkinson, V. Briolat, D. Gent, C. Morrissey, L. A. H. Zeef, G. I. Rice, J.-P. Levraud, and Y. J. Crow. 2015. Characterization of samhd1 morphant zebrafish recapitulates features of the human type i interferonopathy Aicardi-Goutières syndrome. J. Immunol. 194.

64. Guerra, S., L. A. López-Fernández, M. A. García, A. Zaballos, and M. Esteban. 2006. Human gene profiling in response to the active protein kinase, interferon-induced serine/threonine protein kinase (PKR), in infected cells: Involvement of the transcription factor ATF-3 in PKR-induced apoptosis. J. Biol. Chem. 281: 18734–18745.

65. Meng, X., D. Yang, R. Yu, and H. Zhu. 2015. EPSTI1 is involved in IL-28A-mediated inhibition of HCV Infection. Mediators Inflamm. 2015: 716315.

66. Rothenburg, S., N. Deigendesch, K. Dittmar, F. Koch-Nolte, F. Haag, K. Lowenhaupt, and A. Rich. 2005. A PKR-like eukaryotic initiation factor 2 kinase from zebrafish contains Z-DNA binding domains instead of dsRNA binding domains. Proc. Natl. Acad. Sci. 102: 1602–1607.

67. Sepahi, A., L. Tacchi, E. Casadei, F. Takizawa, S. E. LaPatra, and I. Salinas. 2017. CK12a, a CCL19-like Chemokine That Orchestrates both Nasal and Systemic Antiviral Immune Responses in Rainbow Trout. J. Immunol. 199: 3900–3913.

68. Nomiyama, H., N. Osada, and O. Yoshie. 2013. Systematic classification of vertebrate chemokines based on conserved synteny and evolutionary history. Genes to Cells 18: 1–16.

69. Chen, J., Q. Xu, T. Wang, B. Collet, Y. Corripio-Miyar, S. Bird, P. Xie, P. Nie, C. J. Secombes, and J. Zou. 2013. Phylogenetic analysis of vertebrate CXC chemokines reveals novel lineage specific groups in teleost fish. Dev. Comp. Immunol. 41: 137–152.

70. Torraca, V., C. Cui, R. Boland, J.-P. Bebelman, A. M. van der Sar, M. J. Smit, M. Siderius, H. P. Spaink, and A. H. Meijer. 2015. The CXCR3-CXCL11 signaling axis mediates macrophage recruitment and dissemination of mycobacterial infection. Dis. Model. Mech. 7: 253–269.

71. Machala, E. A., S. Avdic, L. Stern, D. M. Zajonc, C. A. Benedict, E. Blyth, D. J. Gottlieb, A. Abendroth, B. P. McSharry, and B. Slobedman. 2018. Restriction of Human Cytomegalovirus Infection by Galectin-9. J. Virol. 93: e01746–18.

72. Abdel-Mohsen, M., L. Chavez, R. Tandon, G. M. Chew, X. Deng, A. Danesh, S. Keating, M. Lanteri, M. L. Samuels, R. Hoh, J. B. Sacha, P. J. Norris, T. Niki, C. M. Shikuma, M. Hirashima, S. G. Deeks, L. C. Ndhlovu, and S. K. Pillai. 2016. Human Galectin-9 Is a Potent Mediator of HIV Transcription and Reactivation. PLoS Pathog. 12: e1005677.

73. Zhu, C., A. C. Anderson, A. Schubart, H. Xiong, J. Imitola, S. J. Khoury, X. X. Zheng, T. B. Strom, and V. K. Kuchroo. 2005. The Tim-3 ligand galectin-9 negatively regulates T helper type 1 immunity. Nat. Immunol. 6: 1245–1252.

74. Brosseau, C., L. Colas, A. Magnan, and S. Brouard. 2018. CD9 tetraspanin: A new pathway for the regulation of inflammation? Front. Immunol. 9: 2316.

75. Richardson, R. B., M. B. Ohlson, J. L. Eitson, A. Kumar, M. B. McDougal, I. N. Boys, K. B. Mar, P. C. De La Cruz-Rivera, C. Douglas, G. Konopka, C. Xing, and J. W. Schoggins. 2018. A CRISPR screen identifies IFI6 as an ER-resident interferon effector that blocks flavivirus replication. Nat. Microbiol. 3: 1214–1223.

76. Cheriyath, V., D. W. Leaman, and E. C. Borden. 2011. Emerging Roles of FAM14 Family Members (G1P3/ISG 6–16 and ISG12/IFI27) in Innate Immunity and Cancer. J. Interf. Cytokine Res. 31: 173–181.

77. Xue, B., D. Yang, J. Wang, Y. Xu, X. Wang, Y. Qin, R. Tian, S. Chen, Q. Xie, N. Liu, and H. Zhu. 2016. ISG12a Restricts Hepatitis C Virus Infection through the Ubiquitination-Dependent Degradation Pathway. J. Virol. 90: 6832–6845.

78. Taylor, H. E., A. K. Khatua, and W. Popik. 2014. The Innate Immune Factor Apolipoprotein L1 Restricts HIV-1 Infection. J. Virol. 88: 592–603.

79. Mekhedov, S. L., K. S. Makarova, and E. V. Koonin. 2017. The complex domain architecture of SAMD9 family proteins, predicted STAND-like NTPases, suggests new links to inflammation and apoptosis. Biol. Direct 12: 13.

80. Meng, X., F. Zhang, B. Yan, C. Si, H. Honda, A. Nagamachi, L. Z. Sun, and Y. Xiang. 2018. A paralogous pair of mammalian host restriction factors form a critical host barrier against poxvirus infection. PLoS Pathog. 14: e1006884.

