## Supplementary material for "Interferon-stimulated genes in zebrafish and human define an ancient arsenal of antiviral immunity": Figure S1

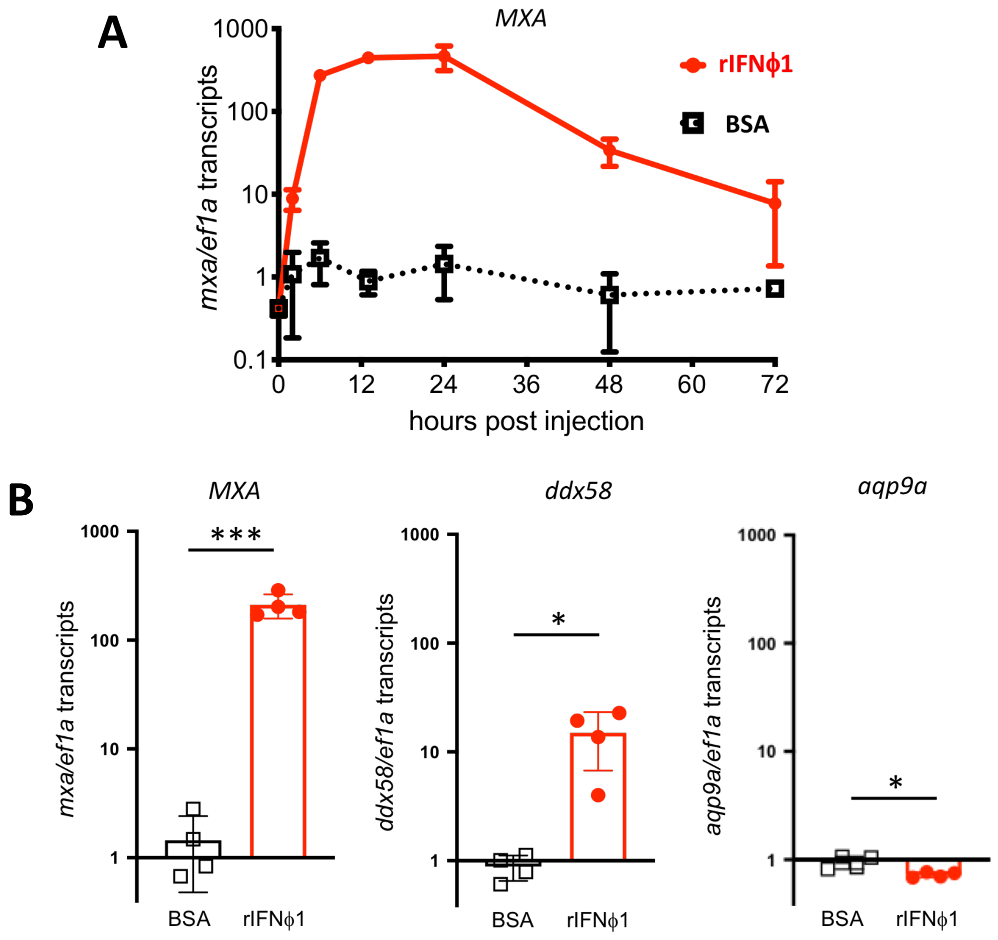


**Figure S1. A. Kinetics of response to recombinant IFNϕ1**, measured by RT-qPCR. Data normalized on the mean of expression of control (samples), displayed as mean ± SD. **B. Genes missing in the reference genome: *ddx58,* encoding RIG-I, is a zebrafish ISG, but *aqp9a* is not.** qRT-PCR in individual zebrafish larvae 6 hours after IFNϕ1 injection; quantification of *MXA* also shown as a control.
