## Supplementary material for "Interferon-stimulated genes in zebrafish and human define an ancient arsenal of antiviral immunity": Figure S2

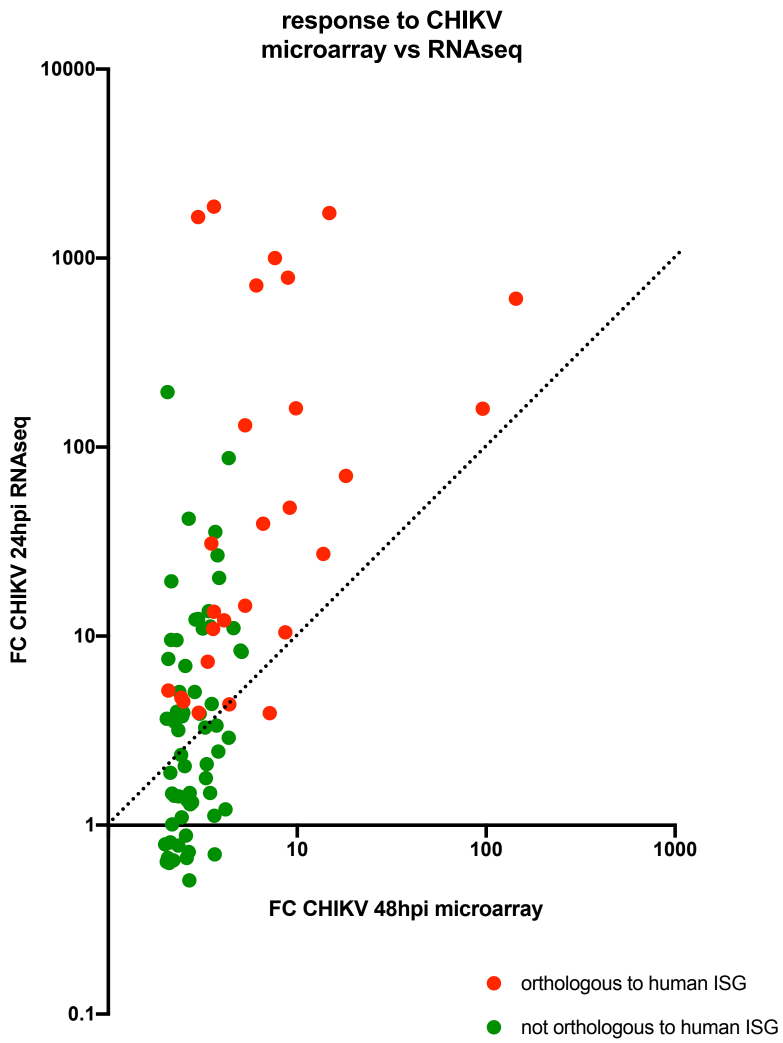


**Figure S2. Transcriptomic response to CHIKV at 24 and 48 hours post infection.** 48h data from (Briolat et al., 2014), 24h data from this study; only genes identified as upregulated in the 2014 microarray study are displayed here.
